# The p21 dependent G2 arrest of the cell cycle in epithelial tubular cells links to the early stage of renal fibrosis

**DOI:** 10.1101/560078

**Authors:** Takayuki Koyano, Masumi Namba, Tomoe Kobayashi, Kyomi Nakakuni, Daisuke Nakano, Masaki Fukushima, Akira Nishiyama, Makoto Matsuyama

**Author notes:** To whom correspondence should be addressed to Makoto Matsuyama; Division of Molecular Genetics, Shigei Medical research Institute, 2117 Yamada, Minami-ku, Okayama, 701-0202, Japan.

## Abstract

Renal fibrosis is accompanied with the progression of chronic kidney disease (CKD). Despite a number of past and ongoing studies, our understanding of the underlying mechanisms remains elusive. Here we explored the progression of renal fibrosis by using a mouse model, unilateral ureter obstruction (UUO). We found that in the initial stage of the progression where extracellular matrix did not deposit yet, the proximal tubular cells arrested at the G2 of the cell cycle. This G2 arrest was induced prior to activation of both DNA damage checkpoint and Wnt/β-Catenin pathway. Further analyses in vivo and in vitro indicated the cyclin dependent kinase inhibitor p21 is involved in the G2 arrest after the damage. The newly produced monoclonal antibody against p21 revealed that the p21 levels were sharply upregulated in response to the damage during the initial stage, but dropped down toward the later stage. To examine the function of p21 in the progression of renal fibrosis, we constructed the novel *p21* deficient mice by *i*-GONAD. Compared with wild-type mice, *p21* deficient mice showed the exacerbation of the fibrosis. Thus we propose that during the initial stage of the fibrosis following the renal damage, tubular cells arrest in the G2 phase depending on p21, thereby safeguarding the kidney functions.

## Introduction

Fibrosis is one of the prominent features of organ disorders where extracellular matrix (ECM) components are excessively accumulated. Although numerous studies sought to uncover the underlying mechanisms in animal models, the molecular details remain unclear. Renal fibrosis is also associated with the progression of the chronic kidney disease (CKD) in which effective treatments have not yet been developed. The myofibroblasts which mainly express ECMs and contribute to the progression of renal fibrosis, are delivered from a variety of origins (1–3). Epithelial-to-mesenchymal transition (EMT) in proximal tubular cells is one of the key processes in the renal fibrosis (4). The previous studies indicate that proximal tubular cells are transformed to interstitial fibroblasts or myofibroblasts by activation of various signaling cascades, *e.g.* the transforming growth factor-β (TGF-β) and Wnt pathway (5–9).

A proliferative response to the damage is the hallmark in acute kidney injury (AKI) (10,11). In principle, cell cycle progression is faithfully moderated by the activation of cyclin dependent kinases (CDKs). CDK inhibitors downregulate the activity of cyclin-CDK complexes via their binding (12). The CDK inhibitor p21 belongs to the Cip/Kip superfamily and plays multiple roles in the regulation of not only cell cycle control but also cellular senescence (13–15). In the kidney, *p21* mRNA is not expressed under the unperturbed condition, but induced after the damage (16). The *p21* lacking mice show opposite phenotypes, amelioration and exacerbation, depending on kinds of the damage (17–19). It is recently reported that *p21* deficient mice alleviate the liver fibrosis because of the elimination of senescent liver stellate cells (20). Thus, the role of p21 in the progression of fibrosis appears complex and its critical role is still under debate.

To maintain the genome integrity from various damages, such as DNA double strand break (DSB), cells also possess the checkpoints which comprise a series of signaling pathways (21,22). DNA damage response (DDR) is one of the protective mechanisms of genome instability. The checkpoint kinase Chk1 which is activated upon DSB, phosphorylates various substrates, thereby inducing G2 arrest until the damages are canceled (21,23). Activation of Chk1 is seen in the process of renal damage in rat (24). In addition, the G2/M arrest is also induced after the renal damage (8,11,25). Under the sustained damaged condition, the G2 arrested cells produce pro-fibrogenic factors, including TGF-β and CTGF (26), by which these factors accelerate the renal fibrosis progression (8,11,26). However, the physiological significance of the cell cycle arrest in response to the damage in the initial stage is unknown.

Here we explored the progression of renal fibrosis by implementing new technical tools, the novel *p21* lacking mice and the newly produced monoclonal antibody. Our data show that in the initial stage of the damage, tubular epithelial cells are arrested in the G2 of the cell cycle in mouse model of renal fibrosis. The G2 arrest is induced prior to DNA damage checkpoint and Wnt/β-Catenin pathway activation, partially depending on CDK inhibitor p21. The *p21* deficient mice exacerbate the progression of the fibrosis. These data uncover a new insight of cell cycle arrest and p21 into the complicated regulations of renal fibrosis.

## Results

### Epithelial tubular cells show a proliferative response after the renal injury

To analyze the step-by-step mechanism(s) of the renal fibrosis, we conducted unilateral ureter obstruction (UUO), which is widely used as a model of renal fibrosis, and collected at the various time points after the injury (Fig. 1A). The number of glomerular was not affected noticeably in this model (Fig. 1A) (6). In the initial stage (∼3d), tubulars were expanded and seemed to be damaged by the obstruction, and in the later stage (3d∼), the area of epithelial cells was decreased and interstitial fibrosis was developed (Fig. 1A). As previously reported, proliferative cells, represented by Ki67 staining, increased after the injury (Fig. 1B and C). It is of note that the number of Ki67 positive cells was decreased in the 7 days-damaged kidney (Fig. 1C). Areas stained with antibody against E-Cadherin, a marker for epithelial cells, were also decreased by obstruction (Fig. 1D). About 40% of Ki67 positive cells were co-stained with E-Cadherin on 3 days after the damage (Fig. 1E). These data suggest that epithelial tubular cells quickly response to the injury and re-entry into the proliferative cycle from the quiescent state.

**Figure 1:**
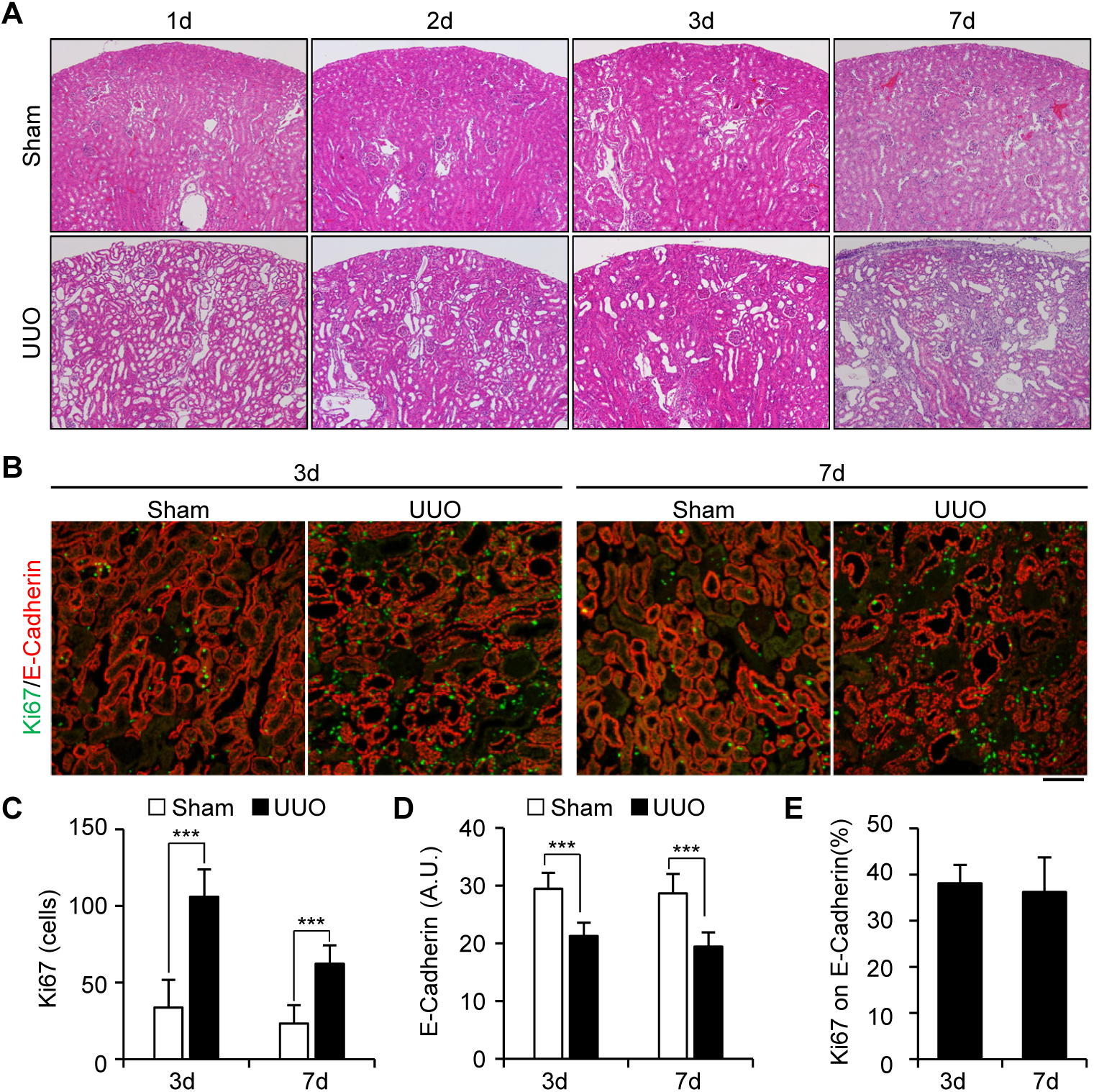
Epithelial tubular cells go into proliferative cycle upon obstruction of kidney. *A*, staining of Hematoxylin and Eosin (HE) of kidney slices obtained from various time points after the obstruction. Sham-operated kidneys were used as a control. The images were taken at x100 magnification. *B*, co-immunostaining with Anti-Ki67 (green) and Anti-E-Cadherin (red). Ki67 was used for recognition of proliferative cells, and E-Cadherin stained the epithelial tubular cells. *Bar*, 10µm. *C* and *D*, quantification of the number of Ki67 positive cells (C) and E-Cadherin stained area (D). *E*, a ratio of Ki67 positive cells on E-Cadherin stained area. The data of obstructed kidneys are shown. ***, *p* < 0.001.

### Epithelial tubular cells arrest at the G2 of the cell cycle prior to renal fibrosis

In order to elucidate the stage of the cell cycle within the damaged epithelial tubular cells, we checked the levels of several cell cycle related proteins. From these experiments, we found that the level of phosphorylated Cdk1 (p-Cdk1^Y15^), which corresponds to an inactivate form Cdk1-cyclinB complex (27), was drastically elevated compared to untreated kidneys during the initial stage of the damage (Fig. 2A). However, this elevation was reduced in the later point, when α-smooth muscle actin (α-SMA), a marker for fibrosis, was accumulated (Fig. 2A). Furthermore, many of p-Cdk1^Y15^ positive cells were detected in the epithelial cells (Fig. 2B). These data suggest that the G2 cells increase within the tubulars in response to the damage.

**Figure 2:**
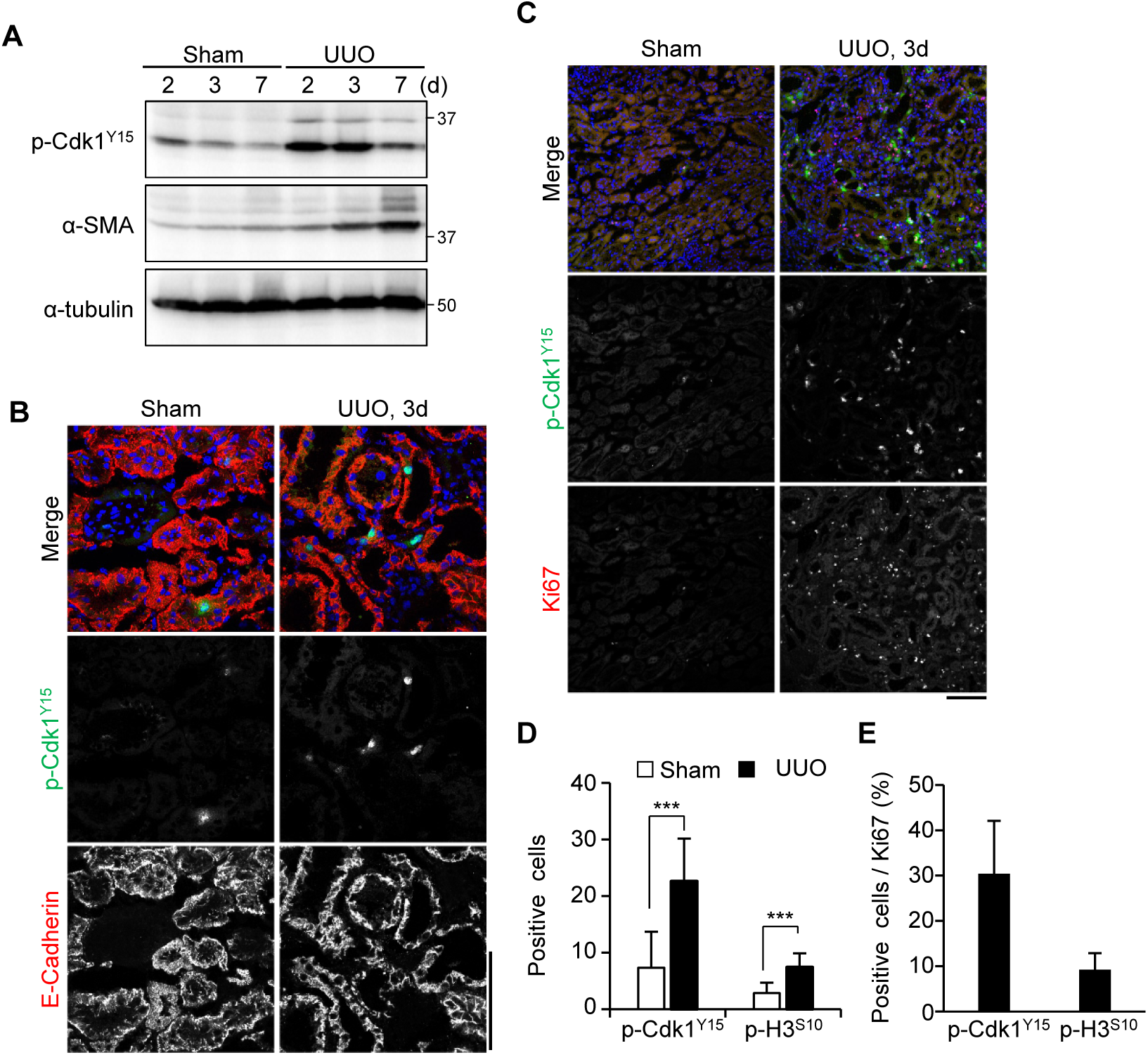
The number of the G2 cells increases prior to development of fibrosis. *A*, immunoblotting of tissue lysates from indicated samples. α-SMA and α-tubulin were used as the marker of fibrosis and the loading control, respectively. The molecular weight (kDa) were shown in the right-hand side of the images. *B*, co-immunostaining with Anti-p-Cdk1^Y15^ (green) and E-Cadherin (red). *C*, co-immunostaining with Anti-p-Cdk1^Y15^ (green) and Anti-Ki67 (red). DAPI (Blue) was used for staining of nucleus. *D*, quantification of the number of p-Cdk1^Y15^ or p-Histone H3^S10^ (p-H3^S10^) positive cells. The quantification was carried out using the sample of 3 days after the obstruction. Each staining were performed using different slices of the same sampling date. *E*, a ratio of p-Cdk1^Y15^ or p-Histone H3^S10^ positive cells merged with Ki67 positive cells. *Bars*, 10μm.

Next, we underwent the double staining of Ki67 and p-Cdk1^Y15^ to measure a population of the G2 cells in the damaged kidneys (Fig. 2C and D). More than 30% of Ki67 positive cells were co-stained with p-Cdk1^Y15^ (Fig. 2E). On the other hand, the positive cells of phosphorylated Histone H3 (p-H3^S10^), which is highly phosphorylated in mitosis (28), were slightly increased (Fig. 2D); however only less than 10% of Ki67 positive cells were merged with p-H3^S10^ (Fig. 2E). These data suggest that epithelial cells arrest prior to mitosis, namely in the G2 phase of the cell cycle, during the initial stage of the damage.

### Tubular cells arrest at the G2 of the cell cycle before activation of DNA damage checkpoint and the Wnt/β-Catenin pathway

To uncover the molecular mechanisms of the G2 arrest upon the renal damage, we firstly examined the relationship with DNA damage checkpoint, because checkpoint activations cause the G2 arrest (21). In addition, an activation of the checkpoint kinase Chk1 after the kidney injury was reported in rat (24). Chk1 activation occurred in a time dependent manner (Fig. 3A). In agreement with this notion, the number of p-H2A.X^S139^ (γH2A.X) which is accumulated onto DNA damaged loci (22), was increased as well (Fig. 3B). This accumulation was detected more frequently when fibrosis was developed (Fig. 3B and C). These data suggest that the activation of DNA damage checkpoint occurs in the later stage of renal damage.

**Figure 3:**
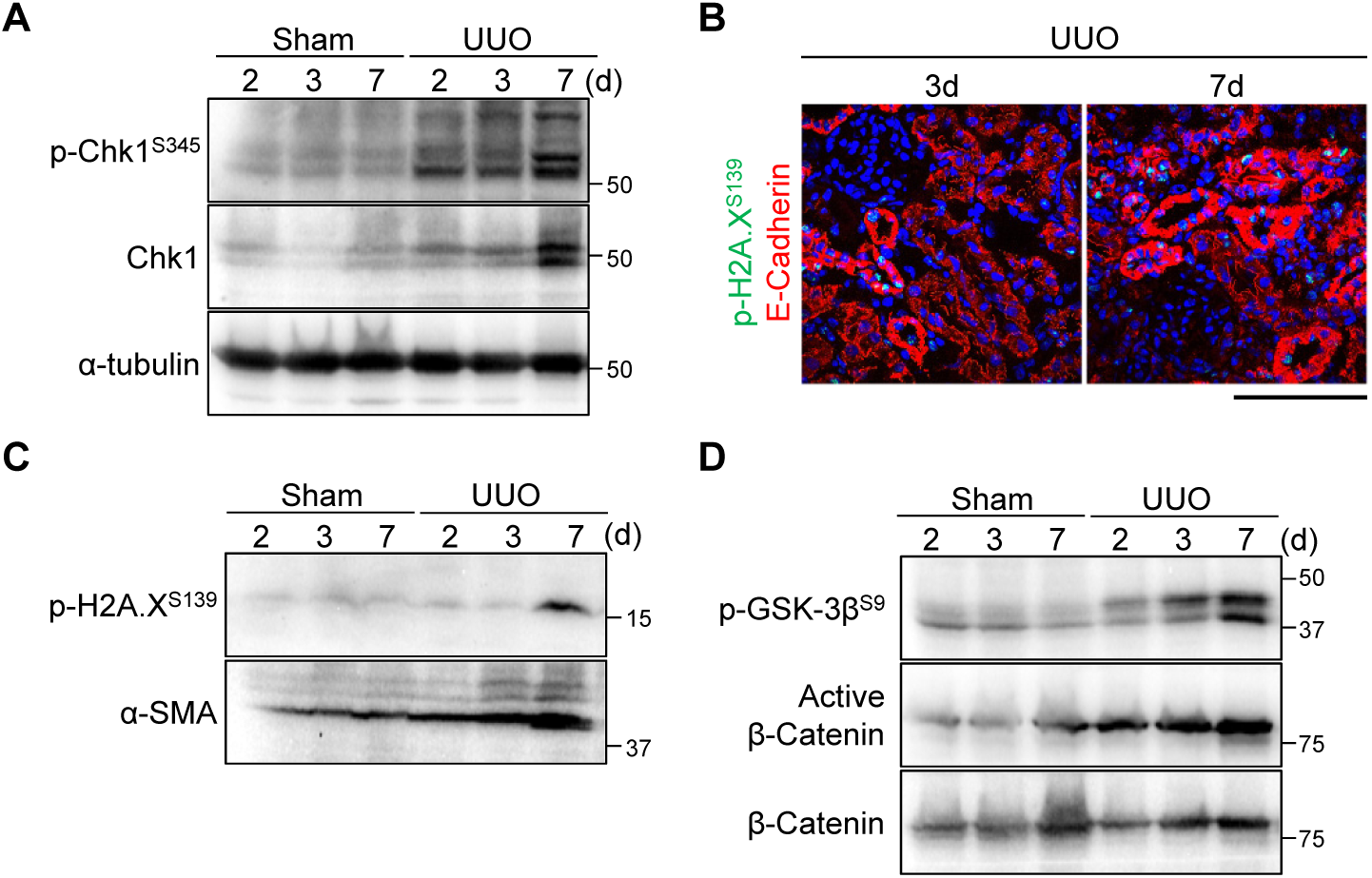
DNA damage checkpoint and Wnt/β-catenin cascade activation are induced in late stage of renal damage. *A*, immunoblotting with indicated antibodies of tissue lysates from indicated samples. *B*, co-immunostaining with Anti-p-H2A.X^S139^ (γH2A.X) (green) and E-Cadherin (red). DAPI (nucleus) was shown in blue. *Bar*, 10µm. *C*, immunoblotting of tissue lysates from indicated samples. α-SMA was used as the marker of fibrosis. Of note, γH2A.X was clearly detected on 7 days after the injury in our hands. *D*, immunoblotting with indicated antibodies of tissue lysates from indicated samples. The molecular weight (kDa) were shown in the right-hand side of the images.

Next, we checked the Wnt/β-Catenin cascade which induces proliferation after the kidney injury (6). Glycogen synthesis kinase-3β (GSK-3β) is implicated in the regulation of Wnt signaling via phosphorylation of β-Catenin (29). Indeed, the phosphorylation of GSK-3β on S9, which is the negative form of GSK-3β, and the active form of β-Catenin (Non-phosphorylated) were increased accompanied with the development of fibrosis (Fig. 3D). Both DNA damage checkpoint and Wnt/β-Catenin pathways were fully activated toward the later stage of renal damage, whereas p-Cdk1^Y15^ level was quickly elevated after the damage (Fig. 2A). Considering these data, we suppose that neither DNA damage checkpoint nor Wnt/β-Catenin is responsible for arrest the G2 phase of the cell cycle in the initial stage of renal damage; instead, these events were induced as a consequence of the G2 cell cycle arrest or redundant regulatory mechanisms.

### p21 is upregulated in response to the initial stage of renal damage

To explore the mechanism of increasing G2 cells, we sought for the responsible molecule(s) by using cultured human tubular epithelial cells (HK-2). To mimic the damage, we added the Aristrochic acid (AA) which induces aristrochic nephropathy (11). DNA damage and the checkpoint activation were induced in a dose dependent manner, mimicking an in vivo damage (Fig. 4A). Intriguingly, p-Cdk1^Y15^ level was elevated in the low concentrations of AA and slightly decreased toward the high concentration even in vitro (Fig. 4A). We found that the expression of CDK inhibitor p21 was also induced by the treatment of AA and the protein level was reduced at the high concentration in vitro (Fig. 4B). Previous work indicates that p21 is upregulated in low doses of DNA damage but transiently degraded in response to high doses of the damage, in which this degradation is induced by phosphorylation carried out by Chk1 (30). Indeed, p21 showed the double bands in Phos-tag containing SDS-PAGE gel, which can detect the phosphorylated protein (Fig. 4B, right-hand side)(31). In addition, the intensity of the band of p21 was decreased at the high concentration of AA (Fig. 4B). These data suggest that p21 is upregulated and degraded by the damage in cultured cells.

**Figure 4:**
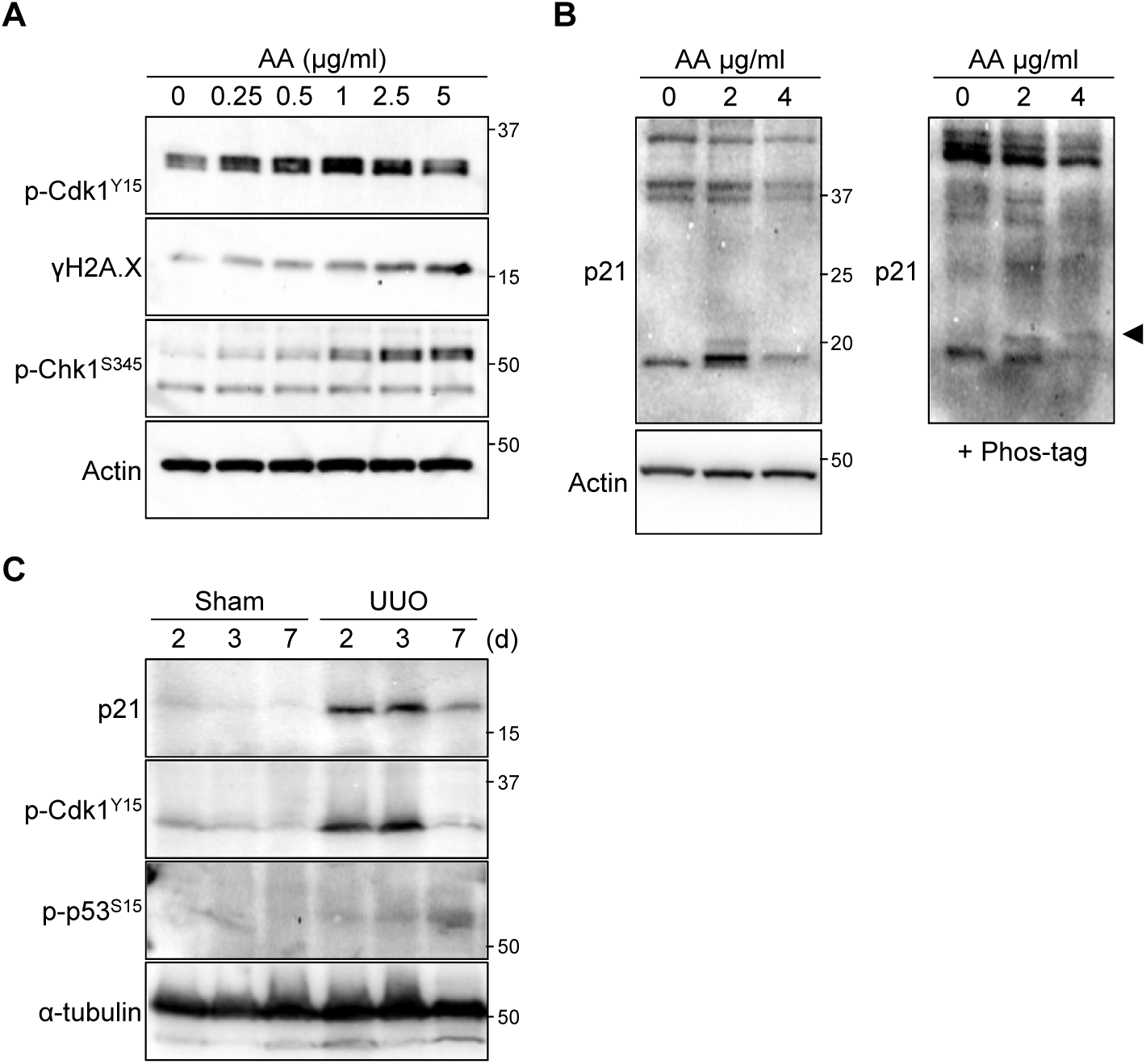
p21 is involved in the response to renal damage. *A*, immunoblotting with indicated antibodies of lysates from Aristrochic Acids (AA) treated HK-2 cells. AA was added into medium, and cells were sampled after 24h later. *B*, immunoblotting with anti-p21. The samples were separated in the presence (right) or absence (left) of 25µM Phos-tag containing SDS-PAGE gel. Arrowhead indicates the phosphorylated band. *C*, immunoblotting with monoclonal p21 antibody of tissue lysates from indicated samples. The molecular weight (kDa) were shown in the right-hand side of the images.

This in vitro data prompted us to examine the relationship between p21 and renal fibrosis. Because the commercial antibodies against p21 were low specificity in mice, we firstly constructed the monoclonal antibody by ourselves to precisely evaluate the protein level of p21 (Fig. S1). The new constructed monoclonal antibody was specific to the mouse p21 in vitro (Fig. S1); and it did not react anything in the *p21* deficient mice (See below). In addition, in healthy kidneys the p21 band reacting with this antibody was not detected (or very low level); by contrast, however, it was sharply upregulated by the obstruction (Fig. 4C). Intriguingly, p21 was also reduced at day 7 damaged kidneys when p-Cdk1^Y15^ level was decreased (Fig. 4C); in other words, the levels of p21 and p-Cdk1^Y15^ displayed a mirror image. The p21 is known as a major target of tumor suppressor p53 and is activated upon the DNA damage (13,14). Although p21 was upregulated during the initial stage of the damage, phosphorylation of p53 on S15 which is induced after the DNA damage (32), was detected only in the later stage (Fig. 4C). These data suggest that p21 is rapidly activated in response to renal damage, though this activation is p53 independent and p21 is then degraded in response to the sustained renal damage.

### Impact of p21 deficiency in renal fibrosis progression

To investigate the impact of p21 in renal fibrosis, we constructed the novel *p21* deficient mice (*p21*^−/−^). To this end, we took the advantages of *i*-GONAD (*i*mproved-*G*enome-editing via *o*viductal *n*ucleic *a*cid *d*elivery) method, in which handling the fertilized egg ex vivo is not required (33,34). Tandem STOP codons were integrated into 60 bases after the first ATG (Fig. S2A). The correct integration was confirmed by both PCR and genome sequence (Fig. S2B and C). Indeed, the p21 protein was not detected even after the UUO in a homozygous deletion (Fig. S2D). These data indicate that the *p21* gene was functionally deleted, similar to a reports previously described (35).

Having constructed *p21*-null mice, we then conducted UUO to both wild-type (WT) and *p21*^−/−^ mice, and sampled at various time points (Fig. 5A). Masson’s trichrome staining revealed that fibrosis is more progressive in *p21*^−/−^ compared to WT mice (Fig. 5B). The staining area of α-SMA was more expanded in *p21*^−/−^ (Fig. 5C and D). These data suggest that global deficiency of *p21* in mice exacerbated the renal fibrosis progression. We underwent further analyses of the renal fibrosis progression in *p21*^−/−^. Double immunostaining of Ki67 and E-Cadherin uncovered that positive cells were increased by *p21* deficiency (Fig. 6A and B). Surprisingly, E-Cadherin measurement showed that more epithelial cells were retained in *p21* deficient mice than WT mice on day 3 after the damage (Fig. 6C). Furthermore, the number of Ki67 positive epithelial cells was not largely changed (Fig. 6D). The immunoblotting showed that the degrees of DNA damage (γH2A.X) and the frequency of apoptotic cells (Cleaved Caspase-3) are not altered either (Fig. 6E). These data suggest that the increased interstitial Ki67 positive cells exacerbate the renal fibrosis progression and that p21 plays some protective role(s) in the epithelial tubular cells in the early stage of the damage.

**Figure 5:**
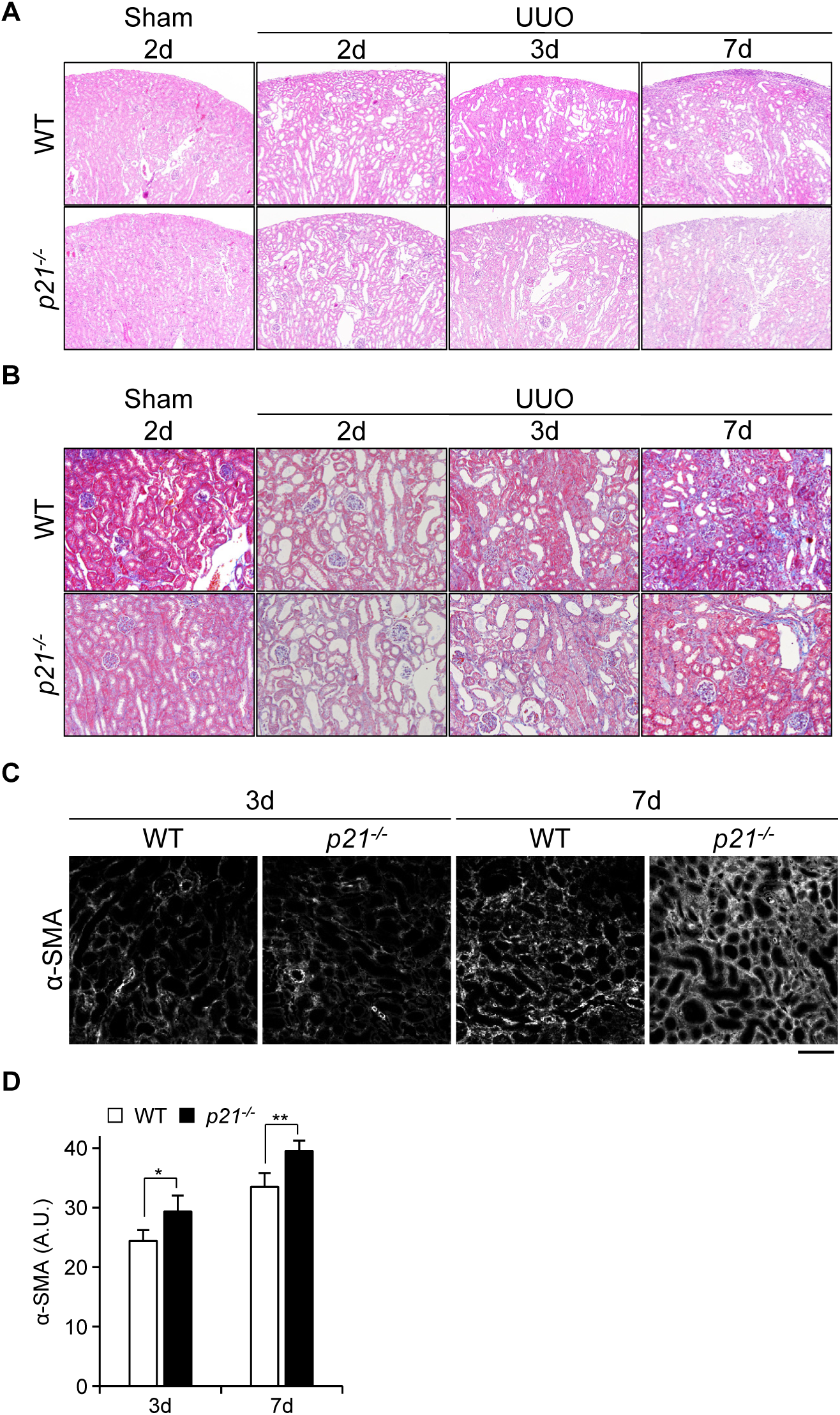
Impact of p21 on renal fibrosis progression. *A* and *B*, histological staining of tissues from WT and *p21*^−/−^ mice. Hematoxylin and Eosin staining (A), and Masson’s trichrome showing fibrosis with blue (B). The images were taken at x100 (A) and at x200 (B) magnification. *C* and *D*, immunostaining staining of obstructed kidneys in WT and *p21*^−/−^ mice. Representative images were shown in C, and quantification was in D. *, *p* < 0.05, **, *p* < 0.01.

**Figure 6:**
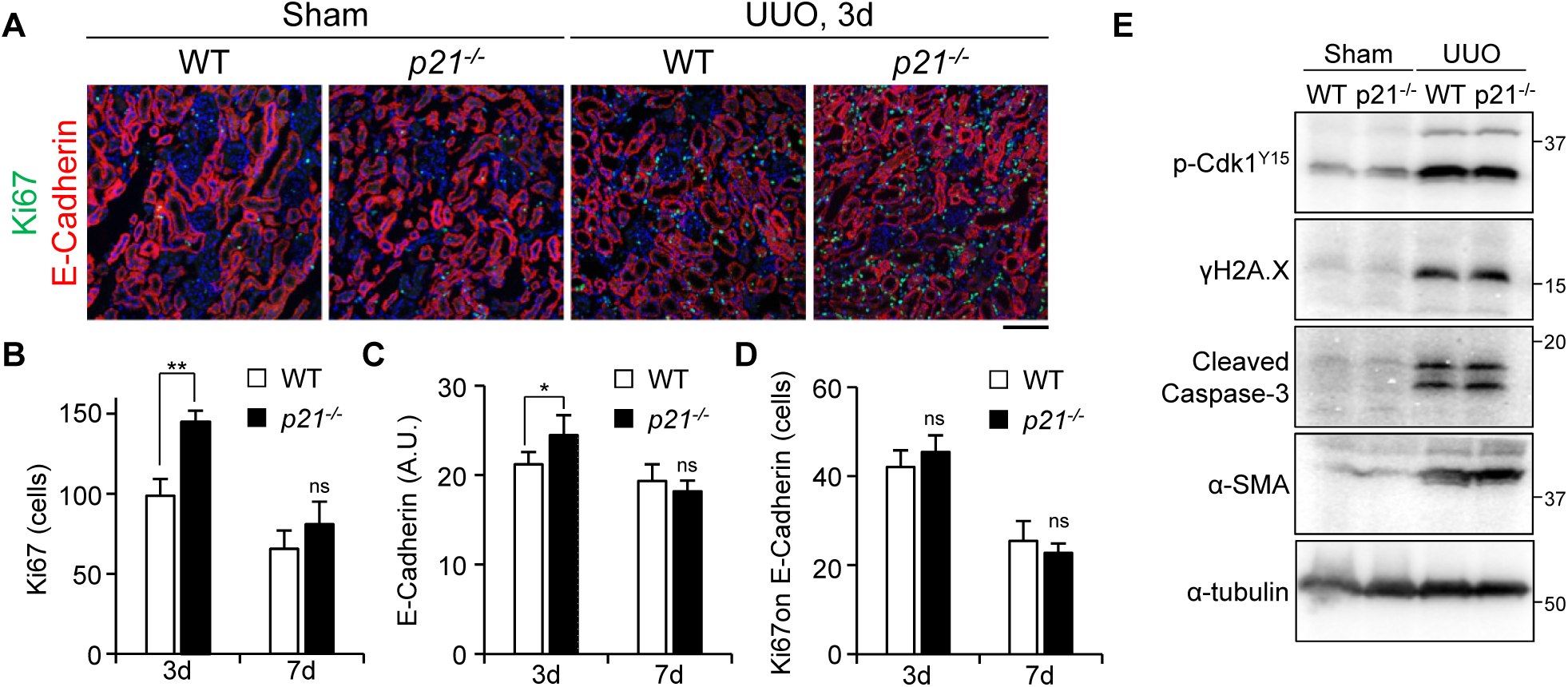
Phenotypes of *p21* deficient mice. *A*, co-immunostaining with Anti-Ki67 (green) and Anti-E-Cadherin (red). DAPI was used for staining of nucleus. *Bar*, 10µm. *B*, quantification of the number of Ki67 positive cells. *C*, quantification of E-Cadherin stained area. *D*, ratio of Ki67 positive cells on E-Cadherin stained area. The data of obstructed kidneys are shown. *E*, immunoblotting with indicted antibodies of tissue lysates from 7days after the injury. *, *p* < 0.05, **, *p* < 0.01.

## Discussion

How fibrosis progresses in kidneys is a fundamental question. Here we addressed this question by implementing two experimental techniques in a mouse model of renal fibrosis: *i*-GONAD to produce the gene modified mice and the monoclonal antibody. Our data showed that the epithelial tubular cells were arrested at the G2 phase of the cell cycle in response to the damage in the initial stage. This G2 cell cycle arrest was partially caused through CDK inhibitor p21. Furthermore, p21 activation and the cell cycle arrest seemed to depend on neither DNA damage checkpoint nor Wnt/β-Catenin. p21 deficiency exacerbated the renal fibrosis compared with wild-type mice. Thus, we propose that the p21 dependent cell cycle arrest contributes to the progression of the renal fibrosis in the early stage of the damage.

### The physiological significance of the G2 cell cycle arrest on renal fibrosis

Our results indicate that the G2 cell cycle arrest is induced even in the early stage of the damage when fibrosis is not developed much yet. Several studies have indicated that the G2 arrest in proximal tubular cells after kidney injury contributes to the progression of the renal fibrosis (8,11,26). We showed that the level of the G2 arrest was inverse proportion to the progression of the fibrosis. The G2 arrest in the initial stage might have a protective role(s) in response to the damage, rather than the production of profibrotic growth factors (26,36). Intriguingly, in the reversible UUO model, the kidney function can be recovered when the obstruction is removed within 2 days (37). In any case, the level of the G2 cell cycle arrest in proximal tubular cells is one of the features in the early stage of renal damage, thus a study for using as a biomarker to predict the fibrosis in initial stage of the damage would be interesting.

### Roles of p21 in the regulation of renal fibrosis

We showed that p21 is involved in the G2 cell cycle arrest in the initial stage of the renal damage. The constructed monoclonal antibody detected the p21 protein in the only damaged kidneys. Unfortunately the p21 monoclonal antibody could not be used for immunostaining. Which cell(s) and/or region(s) express p21 after injury is one of the intriguing questions to be explored. The expression patterning of proteins is very similar to p-Cdk1^Y15^. We suppose that the epithelial tubular cells express p21 and subsequently induce the G2 arrest after the injury to prevent the renal fibrosis progression. Consistent with our finding, the level of *p21* mRNA is increased after the kidney injury even in p53 null mice (16). And we further demonstrated that p21 is degraded in the later stage of the damage when DNA damage is clearly increased. Previous study showed that the regulation of p21 is bimodal, and its level increased in response to the low dose of DNA damage but transiently degraded upon the high level of DNA damage (30). The functions of p21 are regulated by multiple phosphorylation carried by various kinases including the checkpoint kinase Chk1 (13). We speculate that p21 is upregulated in response to the renal damage in the initial stage to protect epithelial tubular cells; however, sustained renal damages cause the DNA damage, leading to Chk1 dependent phosphorylation and degradation of p21.

Previous report indicates that *p21* deficiency ameliorates the development of the fibrosis and the progression of chronic renal failure after the partial renal ablation (17). On the other hand, p21 has a protective and beneficial role(s) in renal ischemia/reperfusion injury (IRI) (18,19). In our study, p21 is initially activated and subsequently degraded in the later stage. And *p21* deficient mice show the severe fibrotic features. It seems that p21 has a protective role(s) in the progression of renal fibrosis. Nonetheless, it is possible that p21 deficiency is beneficial for the fibrosis. In a recent study, *p21* knockout mice improved liver fibrosis because of the elimination of senescent liver stellate cells (20). Our measurement of E-Cadherin dissipation was slightly improved on day 3 in *p21*^−/−^, although the progression of fibrosis was accelerated. This paradoxical functions might be attributed to different cell types and the dose of the damage used. Conditional p21 knockout mice, like cell type specific or drug inducible, will be useful for understanding these issues. Further analyses will illustrate the role(s) of p21 in the regulation of fibrosis.

### The advantage of i-GONAD in the animal research field

Genome manipulation is powerful to understand the molecular mechanisms in the fields of life science and medical research. Recent advances in the genome-editing technology have enabled rapid generation of genome-edited animals with ease. However, the procedure for producing such animals involves multiple complicated steps and also necessitates special techniques, *e.g.* handling zygotes ex vivo. Time consuming process is also one of the limitations when we wish to perform genetic analysis in animals. In this study, we generated the new *p21* deficient mice by using *i*-GONAD method (34), though p21 lacking mice have already existed (35). Our constructed mice showed no production of the protein even after the injury (Fig.4C). Thus, *i*-GONAD method will enable us to analyze the phenotypes of knockout mice rapidly and efficiently in the field of animal research.

## Experimental Procedures

### Mice

C57BL/6 mice were used all in this study. *p21* deficient mice were generated by *i*-GONAD method as previously described (34). In brief, the single strand DNA which contains 3 STOP codons and homology sequence against to the second exon of *Cdkn1a* (cggtcccgtggacagtgagcagttgcgccgtgattgcgattgactagctagaattcccgggcgctcatggcgggctgtctccaggaggcccgagaacggt) was injected to the oviductal lumen of pregnant female mice (E0.7) with Alt-R™ CRISPR-Cas9 system (Integrated DNA Technologies), subsequently electroporated by NEPA21 (Neppa Gene). Guide RNA was designed using CHOPCHOP (http://chopchop.cbu.uib.no/). All materials and equipments were used in accordance with manufacture’s procedures. The correct integration was confirmed by PCR and genome sequence of pups. Renal fibrosis was induced by ligation of left ureters to age matched *p21^+/+^* (WT) and homozygous *p21* deficient mice (*p21*^−/−^), and collected at the indicated time points after the operation (6). Unobstructed kidney (right kidney) were used as controls (sham). All animal experiments were approved by the institutional committee. Animals were handled strictly with the appropriate accordance.

### Antibodies

The rat-monoclonal antibody against to p21 was constructed as previously described (38). Briefly, His-tagged full length of mouse p21 (mp21) was expressed in BL21-CodonPlus-RP. Because His-mp21 was insoluble, we purified the inclusion bodies from the insoluble fraction. Purified His-mp21 with a complete adjuvant was injected into female rats. Other commercial antibodies are listed in table S-1.

### Histological analyses

Kidneys were soaked in 10% buffered neutral formalin at least overnight. Subsequently, the fixed kidneys were embedded in paraffin and sliced with 3µm thickness. The slices were stained with Hematoxylin and Eosin or Masson’s trichrome.

### Immunofluorescence staining

Frozen kidney mounted in compound were sliced with 5µm thickness. The slices were fixed with 10% buffered neutral formalin or 4% paraformaldehyde. After blocking in 5% donkey serum for 30min at room temperature, slices were incubated with primary antibody, followed by Alexa Fluor labeled secondary antibodies (Molecular probe). The images were captured by confocal microscopy (FV2000, Olympus) and were processed by Olympus fluoview ver.4.0. The quantified images were taken by using 20x objective lens (512×512 pixel image) and analyzed by using ImageJ.

### Immunoblotting

Obtained kidneys were homogenized in buffer (50mM Tris, 150mM NaCl, 1mM EDTA, 1% TritonX100) containing with protease inhibitor (Nacalai tesque) on ice and lysates were cleared by centrifuge at 15,000rpm for 15min at 4°C. Cultured HK-2 cells were directly homogenized in a SDS-PAGE sample buffer. Each samples were separated by SDS-PAGE gel and analyzed by western blotting with appropriate antibodies. For the detection of p21 phosphorylation, phosphate affinity SDS-PAGE was carried out by using Phos-tag acrylamide (NARD Institute) (39).

### Cell culture

Human tubular cells HK-2, were purchased from ATCC. HK-2 cells were cultured in keratinocyte serum free medium (K-SFM, Gibco). Cells were treated with indicated concentrations of Aristrochic Acid (AA, Sigma-Aldrich). Dimethyl sulfoxide (DMSO) was used as a solvent.

### Statistical Analysis

Experiments were repeated at least 3 times in each animals. All statistical analyses were carried out using two-tailed unpaired *Student’s* t-test. Each *p*-values were indicated in figure legends.

## Acknowledgements

We are grateful to Drs Takashi Toda and Tohru Okigaki for critical reading of the manuscripts and useful suggestions, to Chieko Takahashi and Mayumi Kohno for supporting animal care. We thank Dr Fumihiro Shigei, the chairman of the board, Sowa-kai Medical Foundation, for encouraging us and financial support. This work was also supported by The KAWASAKI Foundation for Medical Science and Medical Welfare (TK), Wesco Scientific Promotion Foundation (TK, MM), and Ryobi Teien Memory foundation (TK, MM).

## Author contributions

TK and MM designed the project, and TK performed the majority of the experiments and data analyses. TK, MN, TK, and MM constructed and maintained the *p21* deficient mice. KN supported the histological analyses. DN, MF, and AN provided the materials and critical suggestions. TK wrote the manuscript with suggestions from co-authors.

## Abbreviations

CDK: cyclin dependent kinase
UUO: unilateral ureter obstruction
ECM: extracellular matrix
i-GONAD: improved-genome editing via oviductal nucleic acid delivery
CKD: chronic kidney disease
AKI: acute kidney injury
DSB: double strand break
DDR: DNA damage response

## Supporting information

**Table S-1:** Antibodies used in this study.

|  | Company | Clone / Catalog# | Host | Application |
| --- | --- | --- | --- | --- |
| Ki67 | abcam | SP6(ab16667) | Rabbit | IF (1/400) |
| Ki67 | DakoCytomation | TEC3(M7249) | Rat | IF (1/200) |
| E-Cadherin | abcam | DECMA-1(ab11512) | Rat | IF (1/800) |
| p-CDK1<sup>Y15</sup> | abcam | ERP7875(ab133463) | Rabbit | IF (1/200), WB(1/4,000) |
| p-Histone H3<sup>S10</sup> | Cell signaling | D2C8(#3377) | Rabbit | IF (1/200) |
| p-GSK3<sup>S9</sup> | Cell signaling | D85E12(#5558) | Rabbit | WB(1/2,000) |
| p-Chk1<sup>S345</sup> | Cell signaling | 133D3(#2348) | Rabbit | WB(1/1,000) |
| Chk1 | MBL | DCS310(K0086-3) | Mouse | WB(1/1,000) |
| Activated β-Catenin | Cell signaling | D13A1(#8814) | Rabbit | WB(1/2,000) |
| β-Catenin | BD Biosciences | 610153 | Mouse | WB(1/10,000) |
| α-tubulin | Sigma | DM1A(T9026) | Mouse | WB(1/40,000) |
| α-SMA | Cell Signaling | 1A4(#48938) | Mouse | IF (1/500), WB(1/10,000) |
| p-H2A.X<sup>S139</sup> | Cell signaling | 20E3(#9718) | Rabbit | IF (1/500), WB(1/1,000) |
| p21 | Cell signaling | 12D1(#2947) | Rabbit | WB(1/2,000) |
| Cyclin B1 | abcam | ab2949 | Rabbit | WB(1/1,000) |
| Cyclin D1 | abcam | SP4(ab16663) | Rabbit | WB(1/1,000) |
| p-p53<sup>S15</sup> | Cell signaling | D4S1H(#12571) | Rabbit | WB(1/1,000) |
| Cleaved Caspase-3 | Cell signaling | 5A1E(#9664) | Rabbit | WB(1/1,000) |
| Myc-tag | Cell signaling | 9B11(#2276) | Mouse | WB(1/10,000) |
| GFP | Roche | Clones(7.1 and 13.1) | Mouse | WB(1/1,000) |

**Figure S-1:**
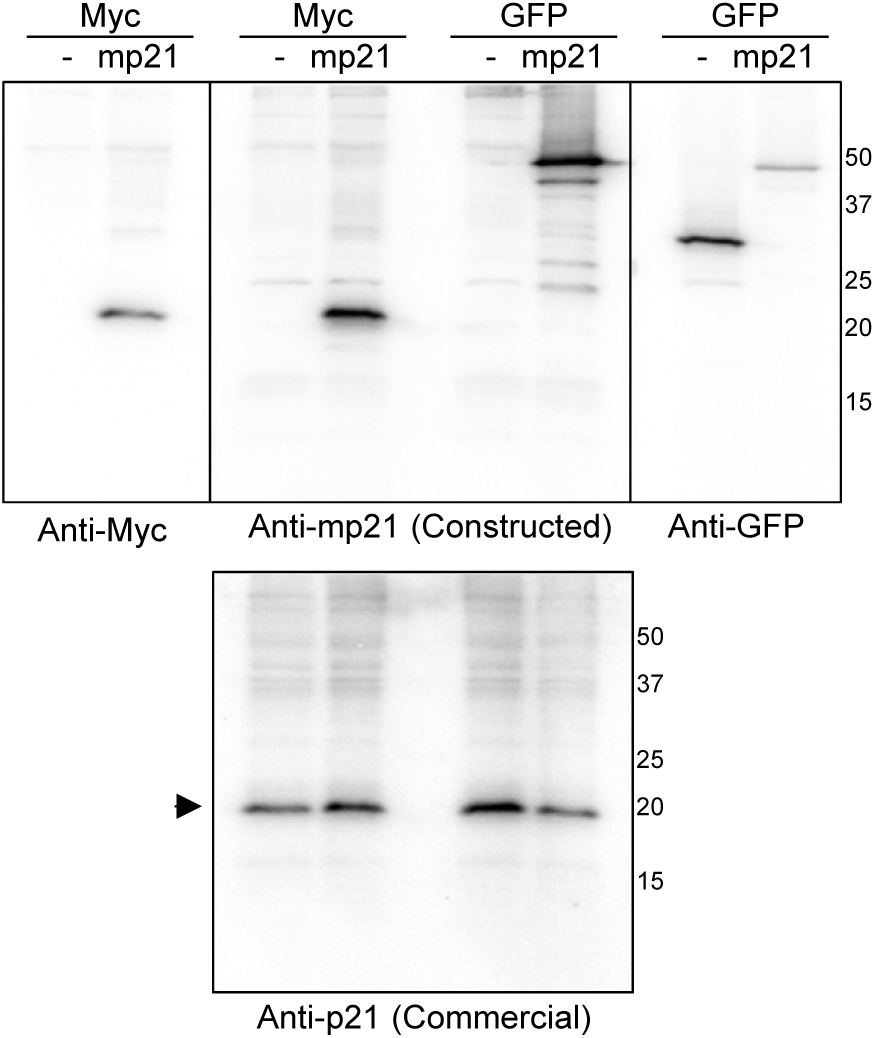
Construction of the monoclonal antibody against mp21. Immunoblotting with the constructed monoclonal antibody. The cell lysates from cos-7 cells expressing Myc-mp21 or GFP-mp21. The indicated antibodies shown in upper panel are used, and the commercial antibody against p21 is used in bottom. Arrowhead indicates the endogenous p21.

**Figure S-2:**
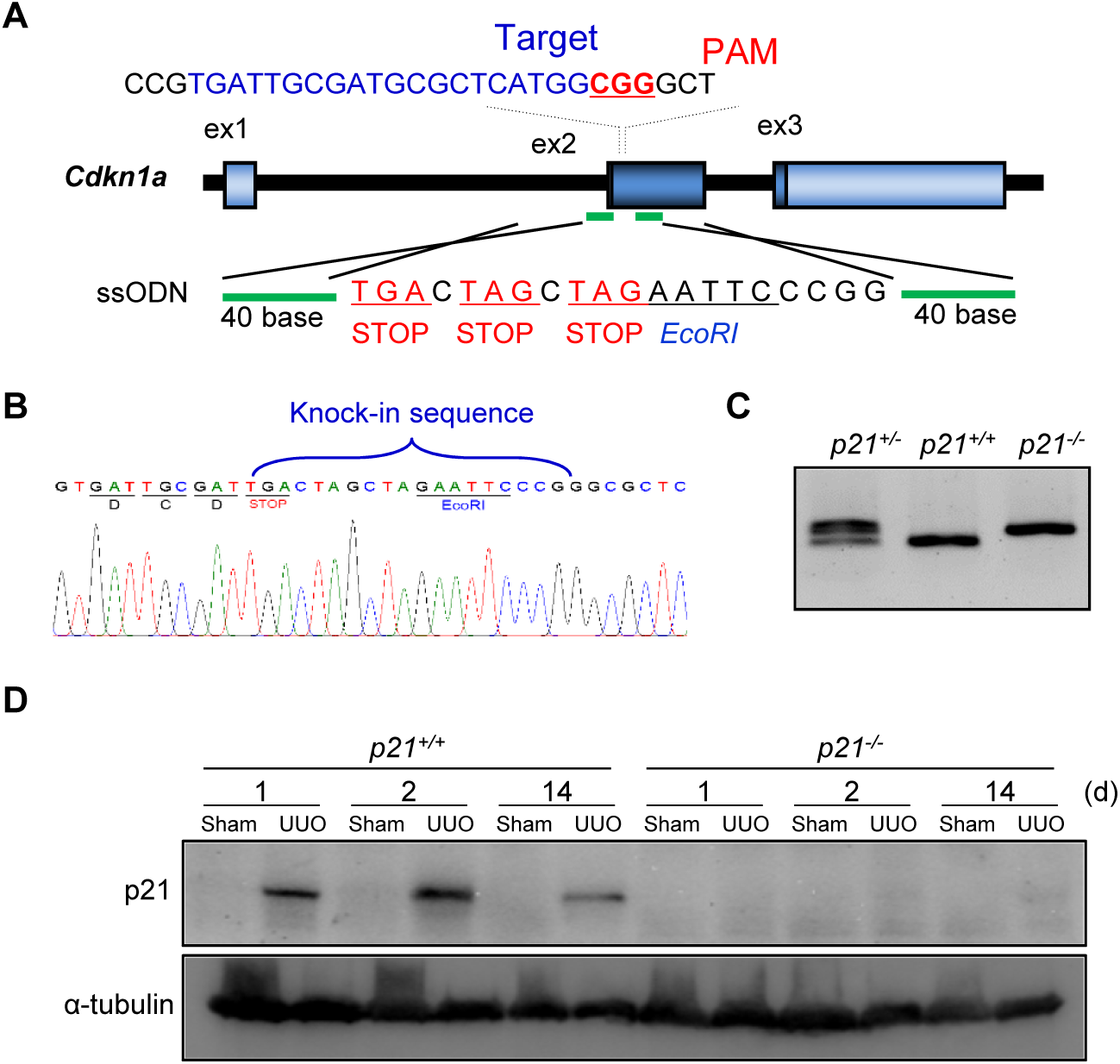
Construction of the *p21* deficiency mice. *A*, the strategy for the construction of *p21* deficiency mice (*p21*^−/−^). *B*, genome sequence of *p21*^−/−^. *C*, the gel image of PCR products amplified from indicated genotypes. *D*, immunoblotting with the mp21 antibody of tissue lysates.

